# A k-mer-based estimator of the substitution rate between repetitive sequences

**DOI:** 10.1101/2025.06.19.660607

**Authors:** Haonan Wu, Antonio Blanca, Paul Medvedev

**Affiliations:** Department of Computer Science and Engineering, The Pennsylvania State University; Department of Biochemistry and Molecular Biology, The Pennsylvania State University; Huck Institutes of the Life Sciences, The Pennsylvania State University

**Keywords:** k-mers, sketching, mutation rates

## Abstract

K-mer-based analysis of genomic data is ubiquitous, but the presence of repetitive k-mers continues to pose problems for the accuracy of many methods. For example, the Mash tool (Ondov et al 2016) can accurately estimate the substitution rate between two low-repetitive sequences from their k-mer sketches; however, it is inaccurate on repetitive sequences such as the centromere of a human chromosome. Follow-up work by Blanca et al. (2021) has attempted to model how mutations affect k-mer sets based on strong assumptions that the sequence is non-repetitive and that mutations do not create spurious k-mer matches. However, the theoretical foundations for extending an estimator like Mash to work in the presence of repeat sequences have been lacking.

In this work, we relax the non-repetitive assumption and propose a novel estimator for the mutation rate. We derive theoretical bounds on our estimator’s bias. Our experiments show that it remains accurate for repetitive genomic sequences, such as the alpha satellite higher order repeats in centromeres. We demonstrate our estimator’s robustness across diverse datasets and various ranges of the substitution rate and k-mer size. Finally, we show how sketching can be used to avoid dealing with large k-mer sets while retaining accuracy. Our software is available at https://github.com/medvedevgroup/Repeat-Aware_Substitution_Rate_Estimator.

## 1 Introduction

*K*-mer-based analysis of genomic data is ubiquitous. e.g. in genome assembly [1], error correction [2], read mapping [13], variant calling [29], genotyping [30, 7], database search [14, 9], metagenomic sequence comparison [26], and alignment-free sequence comparison [28, 20, 24]. One of the major challenges is the presence of repetitive *k*-mers, which adversely affects the practical performance as well as the theoretical analysis of downstream algorithms. One example is that heuristic read aligners like minimap2 [15] and even more rigorous ones like Eskemap [25] filter out highly repetitive *k*-mers in order to avoid explosive run times. Another example is the recent paper [27] that proved that sequence alignment can on average be done in almost 𝒪 (*n* log *n*) time but could not account for sequences with a high number of repeats.

One of the major advantage of *k*-mer-based methods is that they lend themselves more easily to sketching [16, 22], which is important for scaling to large-scale data. The ground-breaking Mash paper [20] was able to estimate the mutation rate between two genomes fast enough to be able to construct a phylogeny of 17 primate species in a tiny fraction of the time it would take an alignment-based method. Their approach uses an estimator based on the Jaccard similarity between the *k*-mer sketches of two sequences. However, the derivation behind their estimator assumes that the genomes have no repeats, making it inaccurate in highly repetitive regions. Other methods for estimating mutation rates are not designed for and/or not tested on highly repetitive sequences [34, 10, 18, 23].

In this paper we tackle the challenge of accounting for repeats when estimating the mutation rate. We assume that a string *t* is generated from a string *s* through a simple substitution process [5], where every nucleotide of *s* mutates with a fixed probability *r*. Given the number of shared *k*-mers between *s* and *t* and the *k*-mer abundance histogram of *s*, we define our estimator 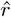 as the solution to a polynomial equation, which can be solved using Newton’s method. We give a theorem to bound its bias, in terms of properties of *s* (Theorem 3). Our estimator is designed to capture the most salient properties of the repeat structure of the genome, with the rest of the information being captured in the bias bounds. As a result, a user can decide *a priori* whether to trust our estimator, based on the quality of the bias bounds and on another heuristic we provide (Theorem 4).

We evaluate our estimator 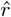 empirically on various sequences, including the alpha satellite centromeric region of human chr21 and the highly repetitive human RBMY gene. For such repetitive sequences, our estimator remains highly accurate, while the repeat-oblivious estimator of the kind used by Mash is unreliable. We make a comprehensive evaluation of 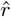 across the spectrum of *k* and *r* values, which can guide a user towards choosing a *k* value for their analysis. We also show that our estimator can be used on top of a FracMinHash sketch, without systematically effecting the bias. Our software is available on GitHub [32].

## 2 Preliminaries

Let *s* be a string and let *k >* 0 be a parameter indicating the *k*-mer size. We will index string positions from 1. We further assume in this paper that *s* is at least *k* nucleotides long. We use *L* to denote the number of nucleotides in the string minus (*k* − 1), or, equivalently, the number of *k*-mers in *s*. For 1 *≤ i ≤ L*, let *s*_*i*_ be the *k*-mer starting at position *i* of *s*. Let *sp*^*k*^(*s*) be the set of all distinct *k*-mers in *s*, also called the *k-spectrum* of *s*. We let *L*_0_ be the size of *sp*^*k*^(*s*), i.e. the number of distinct *k*-mers in *s*. Given a *k*-mer *τ*, we will use the shorthand *τ* ∈ *s* to mean *τ* ∈ *sp*^*k*^(*s*). Given two strings *s* and *t*, we define *I*(*s, t*) ≜ |*sp*^*k*^(*s*) ∩ *sp*^*k*^(*t*)| as the number of *k*-mers shared between them. We will usually use the shorthand of *I* for *I*(*s, t*). Given two *k*-mers *τ* and *ν*, we use HD(*τ, ν*) to denote their Hamming distance.

Let *K* be a set of *k*-mers and let *s* be a string. We let *occ*(*K*) denote the number of positions *i* in *s* such that *s*_*i*_ ∈ *K*. When *K* consists of a single element *τ*, we simply write *occ*(*τ*). A set of positions 𝒥 is said to be a *set of occurrences* of *K* if for all *i* ∈ 𝒥, we have *s*_*i*_ ∈ *K*. A set of occurrences is said to be *non-overlapping* if, for all distinct *i, j* ∈ 𝒥, |*j* − *i*| *≥ k*. We let *sep*(*K*) be the maximum size of a set of non-overlapping occurrences of *K*, also referred to as the *separated occurrence count*. Observe that 0 *≤ sep*(*K*) *≤ occ*(*K*). The *abundance histogram* of a string *s* is the sequence (*a*_1_, …, *a*_*L*_) where *a*_*i*_ is the number of *k*-mers in *sp*^*k*^(*s*) that occur *i* times in *s*. Note that 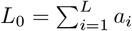.

We will consider the following random *substitution process*, parameterized by a rate 0 *≤ r ≤* 1. Given a string *s*, it generates an equal-length string where, independently, the character at each position is unchanged from *s* with probability 1 − *r* and changed to one of the three other nucleotides with probability *r/*3.

## 3 Problem overview and proposed solution

In this paper, we address the following problem. Let 0 *≤ r ≤* 1 be a substitution rate. Let *s* be a string and let *t* be generated from *s* using the substitution process parametrized by *r*. Let *I*_obs_ = *I*(*s, t*) be the observed spectrum intersection size. Given *I*_obs_ and the abundance histogram of *s*, the problem is to estimate the mutation rate *r*.

The *bias* of an estimator 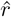 for *r* is defined as 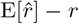. A good estimator should have a small absolute bias, one that is within the error tolerance of downstream applications. For our problem, directly finding an estimator for *r* with provably small bias turned out to be technically challenging. Instead, we provide an estimator for *q* ≜ 1 − (1− *r*)^*k*^, which corresponds to the probability that a *k*-mer occurrence contains at least one substitution. There is a natural one-to-one correspondence between an estimator 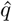 of *q* and an estimator 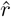 of *r* via the equation 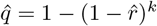. Thus, an alternative to bounding the bias of 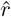 is to bound that of 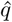; i.e., bound 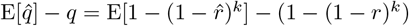. While the difference between the two approaches may intuitively seem minor, bounding the bias of 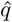 turned out to be more technically feasible.

The only previously known estimator for this problem is what we refer to as the *repeat-oblivious* estimator^1^ and denote by *r*_obl_. The derivation of this estimator assumes that 1) *s* has no repeats and 2) if a *k*-mer mutates, it never mutates to anything that is already in *s*. The estimator is then derived using a simple two step technique, called the method of moments [6]. The first step is to derive E[*I*] in terms of *r*. Under the assumptions (1) and (2), E[*I*] = *L*(1 − *r*)^*k*^. The second step is to take the observed value *I*_obs_, plug it in place of E[*I*], and solve for *r*. In this case, one solves the equation *I*_obs_ = *L*(1 − *r*)^*k*^ and gets the estimator 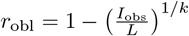 and the corresponding estimator 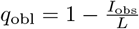. The estimator *q*_obl_ is an unbiased estimator of *q* when (1) and (2) hold, but the bias for general sequences is not known. We are also not aware of any results about the bias of *r*_obl_, even under assumptions (1) and (2). As we will show in Section 6, the repeat-oblivious estimator has a large empirical bias when the assumptions are substantially violated.

On the one hand, the repeat-oblivious estimator does not at all account for the repeat structure of *s*. On the other hand, an estimator that would fully account for it seems to be challenging to derive, analyze, and compute. Moreover, such an estimator would likely be superfluous for real data. Instead, our approach is intended to achieve a middle ground between accuracy and complexity by accounting for the most essential part of the repeat structure in the estimator and expressing the non-captured structure in the bias formula. We will show that under assumption (2) and the assumption that all *k*-mer occurrences are non-overlapping in *s*,

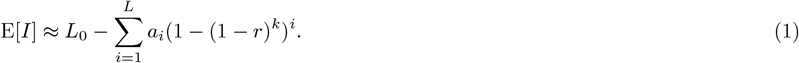

Following the method of moments approach, we define our estimator 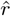 as the unique solution (the uniqueness is shown in Lemma A.2) to this equation when plugging in *I*_obs_ in place of E[*I*]. Though we are not able to analytically solve for 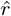, we can find the solution numerically using Newton’s Method.

Note that the assumptions we make are not necessary to compute 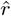 and only represent the ideal condition for our estimator. Our theoretical and experimental results will quantify more precisely how the deviation from our assumptions is reflected in the bias.

## 4 Estimator bias

Recall that we define 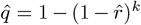 and, as mentioned earlier, we will prove the theoretical results on the bias of 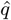, rather than 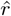. First, we need to derive the expectation and variance of the intersection size. A closed-form expression for even the expectation is elusive, so we will instead use an approximation and derive bounds on the error. The idea behind our bounds is that the error becomes small on the types of sequences that occur in biological data.

We want to underscore that when we make probabilistic statements, it is with regard to the substitution process and not with regard to *s*. We do not make any assumptions about *s*, and, in particular, we are not considering the situation where *s* itself is generated randomly.

First, it is useful to express *I* ≜ *I*(*s, t*) as a sum of indicator random variables. Let us define 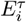 as event that *t*_*i*_ = *τ* and 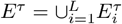 as the event that at least one position in *t* contains *τ*. By linearity of expectation, we have

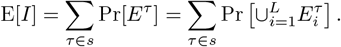

Let 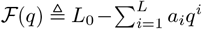. Relying on the approximation E[*I*] *≈* ℱ (*q*) (i.e. Equation (1)), we define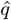 as the solution to *I*_obs_ = ℱ (*q*), or, equivalently, 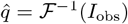. We show that this approximation holds when we assume that 1) 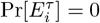 when *s*_*i*_≠ *τ* (see footnote^2^) and 2) all occurrences of *τ* are non-overlapping in *s*:

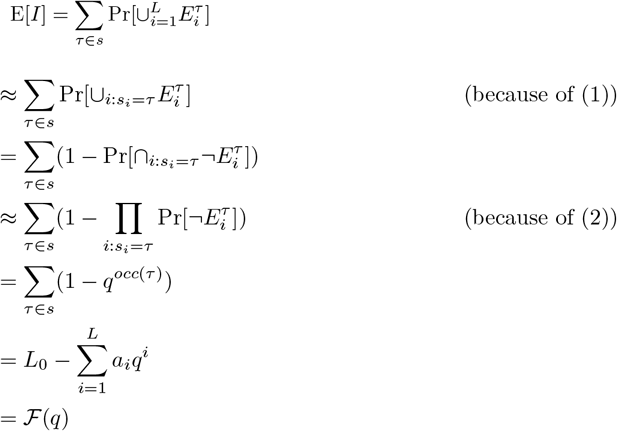

The underlying philosophy for our estimator is that while these assumptions are not perfectly satisfied on real data, in most cases the contribution due to violations of these assumptions is small. To make this mathematically precise, we will bound the difference between E[*I*] and ℱ (*q*) in terms of an expression that can be calculated for any *s*.

### Theorem 1.

*We have that L*_*E*_ *≤* E[*I*] *≤ U*_*E*_, *where*

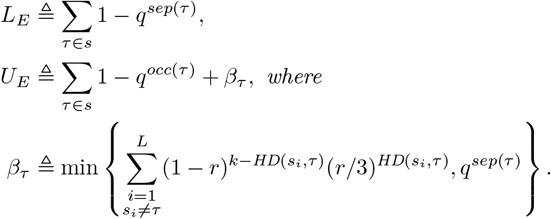

The difference between ℱ (*q*) and *L*_*E*_ (i.e. ∑ _*τ∈s*_ *q*^*sep*(*τ*)^ − *q*^*occ*(*τ*)^) is close to 0 when the number of *k*-mers with overlapping occurrences is close to 0. On the other hand, the difference between ℱ(*q*) and *U*_*E*_ (i.e. 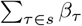) is never zero (except in corner cases). However, the largest terms contributing to this difference are due to pairs of non-identical *k*-mers that have a small Hamming distance to each other. Thus, the difference becomes small when the number of “near-repeats” is small.

Next, we upper bound the variance.

### Theorem 2.

*We have that*

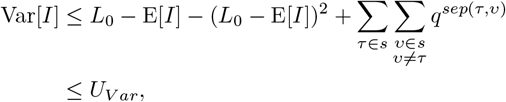

*where*

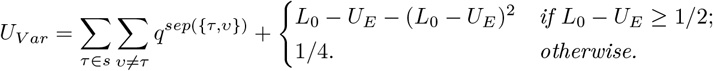

Theorem 2 gives an upper bound on Var[*I*] in two forms. The first one is more precise because it is a function of E[*I*]. However, since we are not able to compute E[*I*] exactly, the second form allows us to plug in the upper bound on E[*I*] from Theorem 1.

Given the bounds on E[*I*] and Var[*I*], we are now able to bound the bias of 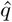.

### Theorem 3.

*Let s be a sequence with at least one k-mer that occurs exactly once. The bias of* 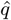 *is* 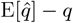 *where* 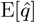 *is bounded as*

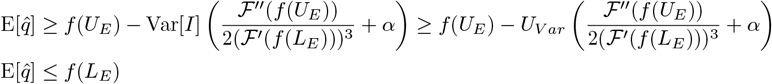

*and where f* ≜ ℱ^−1^ *and* 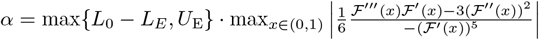.

The derivatives of ℱ (*q*) have straightforward closed-form expressions, since ℱ is a polynomial in *q*. We do not have a closed-form solution for *f*, but it can be evaluated numerically using Newton’s method. Thus, for any given sequence *s*, we can precompute the bounds of our 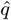 estimator bias for any value of *q*. Due to space limitations, we do not further elaborate on the algorithm to compute the bounds in Theorem 3 or on its runtime analysis.

When the observed intersection is empty, there is a loss of signal and it becomes challenging for any intersection-based estimator to differentiate the true substitution rate from 100%. The following theorem gives an upper bound on the probability that the intersection is empty, as a function of *L, k*, and *r*. In Section 6, we will show how it can be used to make a conservative decision that the computed estimate is unreliable.

### Theorem 4.

*Let s be a string of length at least k. The probability that every interval of length k in s*[1..*i* + *k* − 1] *has at least one substitution can be computed in* Θ(*ik*) *time with a dynamic programming algorithm that takes as input only L, r, k (not s)*.

## 5 Proofs

This section contains the proofs of our theoretical results. In particular, we will prove Theorems 1–3 from the previous section. The proof of Theorem 4 is left for the Appendix. We start by proving a couple of preliminary facts that will be used in the proofs of these theorems. First, we consider the probability of the event 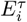, which is straightforward to derive.

### Lemma 5.

*For all* 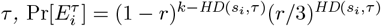.

**Proof**. In order for *s*_*i*_ to be equal to *τ* after the mutation process, exactly *k* − HD(*s*_*i*_, *τ*) positions must remain unmutated (which happens with probability (1 − *r*)^*k*−*d*^) and exactly HD(*s*_*i*_, *τ*) positions must mutate to the needed nucleotide (which happens with probability 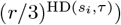. ▪

Next, we will bound the probability that all the occurrences of a *k*-mer become mutated; i.e. a *k*-mer does not survive the mutation process.

### Lemma 6.

*Let τ be a k-mer with occurrence locations denoted by p*_1_ *<* … *< p*_*occ*(*τ*)_. *For all* 2 *≤ ℓ ≤ occ*(*τ*),

1. 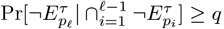, *and*
2. 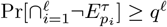.

**Proof**. We drop *τ* from the notation since it remains constant throughout the proof. We first prove the first statement of the lemma. Let us consider the intervals associated with 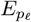 and 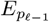, denoted by [*p*_*ℓ*_, *p*_*ℓ*_ + *k* − 1] and [*p*_*ℓ* − 1_, *p*_*ℓ* − 1_ + *k* − 1], respectively. If these intervals are disjoint, then we are done. Otherwise, the union of these intervals can be partitioned into three regions: 1) the part of the interval of 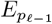 that does not intersect with the interval of *E*_*p*_, 2) the intersection of the two intervals, and 3) the part of the interval of 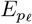 that does not intersect with the interval of *E*_*p ℓ* − 1_. We denote the lengths of these intervals as *a, b*, and *c*, respectively, and we denote the event that no mutation occurs in the intervals as *A, B*, and *C*, respectively. Let 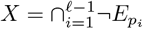, i.e. we need to calculate 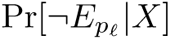. First, we reduce the calculation to Pr[*B*|*X*] as follows:

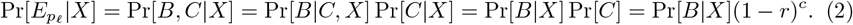

Next, to calculate Pr[*B*|*X*], we proceed by conditioning on *A*:

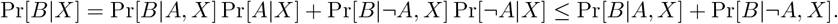

First note that

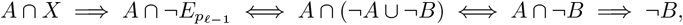

and so Pr[*B*|*A, X*] = 0. To bound Pr[*B*|¬*A, X*], consider all the intervals 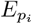, for *i < ℓ*, that intersect with *B*’s interval. Formally, let 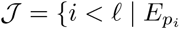 intersects *B*’s interval}. Note that all intervals indexed by 𝒥 necessarily contain *A*’s interval. Therefore, the event ¬*A* implies 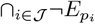. We can now write

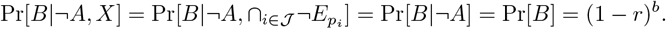

Therefore, Pr[*B* | *X*] *≤* (1 − *r*)^*b*^ and plugging this bound into Equation (2), we get the first statement of the lemma.

To prove the second statement of the lemma, we apply the chain rule together with the first statement:

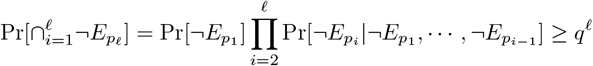

▪

We can now prove Theorem 1:

**Proof of Theorem 1**. It suffices to prove that for every k-mer *τ* ∈ *s*, it holds that 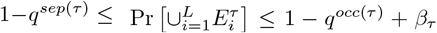. For the lower bound, let 𝒥 be a non-overlapping set of occurrences of *τ* of size *sep*(*τ*). Then we have

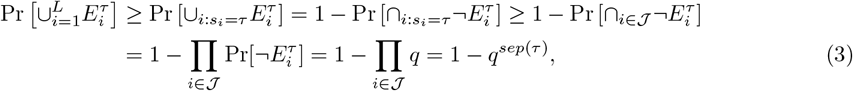

where we use the independence of the events 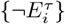 when they are non-overlapping. For the upper bound, let 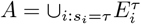 and let 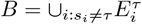. Then, by Lemma 6,

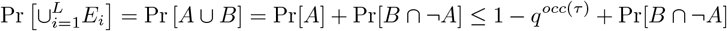

To bound Pr[*B* ∩ ¬*A*] observe that Pr[*B* ∩ ¬*A*] *≤* min(Pr[*B*], Pr[¬*A*]), and by Lemma 5:

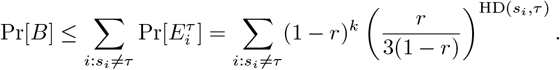

Moreover, by Equation (3), Pr[¬*A*] = 1 − Pr[*A*] *≤ q*^*sep*(*τ*)^ and the result follows. ▪

The proof of the variance bound is more straightforward:

**Proof of Theorem 2**. Since *I* is a sum of indicator random variables (i.e. *I* = ∑_*τ∈s*_ *E*^*τ*^), we can write the variance as

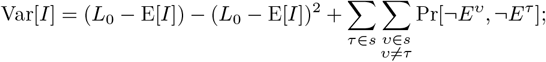

for completeness we include a proof of this fact in the appendix (Lemma A.1).

Consider some *τ*≠ *ν* and let 𝒥 be a non-overlapping set of occurrences of {*τ, ν*}. Let 𝒥 ^*τ*^ *⊆* 𝒥 be the positions where *τ* occurs and let 𝒥 ^*ν*^ *⊆* 𝒥 be the positions where *ν* occurs. Then,

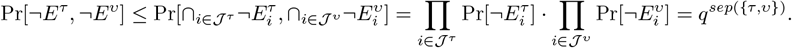

This gives the first form of the upper bound on the variance. The *U*_*V ar*_ upper bound is derived from the fact that *f* (*x*) = *x*−*x*^2^ is monotonically increasing on [0, 1*/*2) and decreasing on [1*/*2, *∞*). Therefore, the maximum of 1*/*4 is achieved at *x* = 1*/*2. ▪

**Proof of Theorem 3**. In Lemma A.2 in the Appendix, we show that *f* is well-defined. We will only consider *f* on the interval [ℱ (1), ℱ (0)]. Throughout the proof, we will rely on the facts that 1) on the interval *q* ∈ [0, 1], ℱ ^*′*^(*q*) *<* 0, ℱ^*′′*^(*q*) *≤* 0, ℱ ^*′′′*^(*q*) *≤* 0; 2) for *y* ∈ [ℱ (1), ℱ (0)], *f*^*′*^(*y*) *<* 0 and *f*^*′′*^(*y*) *≤* 0; 3) the first three derivatives of *f* can be expressed in terms of *f* and the derivatives of ℱ. These properties follow by basic calculus and are stated formally in Lemma A.2. Recall that 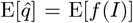. To get the upper bound, we use the fact that *f* is decreasing and concave. We apply Jensen’s inequality followed by Theorem 1 to get that E[*f* (*I*)] *≤ f* (E[*I*]) *≤ f* (*L*_*E*_).

For the lower bound, since we cannot analytically derive *f* (*I*), we derive a reverse Jensen inequality using the Taylor expansion of *f* around E[*I*]. Specifically, using the Lagrange Remainder, we know that there exists some *ξ*_*I*_ between *I* and E[*I*] such that

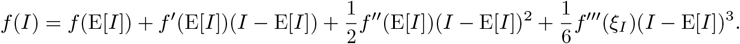

Since we are interested in the expected value of *f* (*I*), we take expectations on both sides:

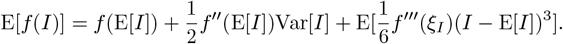

We will bound the terms separately by writing E[*f* (*I*)] *≥ T*_1_ + *T*_2_ − *T*_3_ *·* max_*y*∈[*F* (1),*F* (0)]_ *T*_4_ with 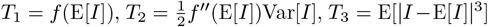, and 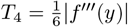. For the first term, we use the fact that *f* is decreasing and apply Theorem 1 to get that *f* (E[*I*]) *≥ f* (*U*_*E*_). For the second term *T*_2_, we first plug in the second derivative of *f* and then apply monotonicity properties together with Theorem 1 to get

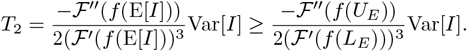

For *T*_3_, we use the fact that *I ≤ L*_0_, which implies that |*I* − E[*I*] |*≤* max(*L*_0_ − E[*I*], E[*I*]), and thus

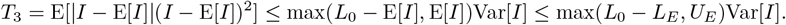

For *T*_4_,

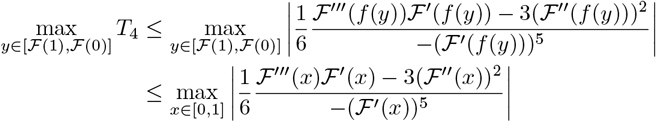

▪

## 6 Experimental results

In this section, we evaluate the empirical accuracy and robustness of our estimator, used by itself or in combination with sketching.

### 6.1 Datasets

To evaluate our estimator, we use four sequences to capture various degrees of repetitiveness. The sequences are extracted from the human T2T-CHM13v2.0 reference [19]. The sequences and their coordinates are available at our reproducibility GitHub page [33]. Table 1 shows properties of these sequences and Figure 1 shows their cumulative abundance histograms.

**Table 1.** Sequence properties of our four experimental datasets. A *k*-mer *τ* is *overlapping* if it overlaps itself at least once in the sequence, i.e. *sep*(*τ*) *< occ*(*τ*).

| Name | Default $k$ | N. $k$-mers<br>($L$) | N. distinct<br>$k$-mers ($L_0$) | N. of overlapping<br>$k$-mers | Biological<br>significance |
| --- | --- | --- | --- | --- | --- |
| *D-easy* | 20 | 100,000 | 98,786 | 15 | arbitrary region |
| *D-med* | 20 | 14,400 | 13,727 | 2 | RBMYA1 gene |
| *D-hard* | 10 | 2,264 | 1,199 | 0 | simple repeat |
| *D-hardest* | 30 | 100,000 | 3,987 | 0 | centromere |

**Figure 1.**
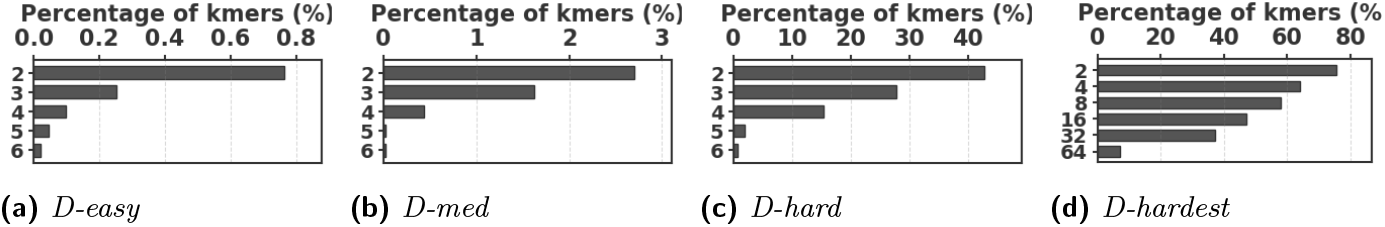
Cumulative abundance histograms of our datasets. Each row labeled *y* shows the percentage of *k*-mers which occur at least *y* times.

1. *D-easy*: This is an arbitrarily chosen substring from chr6, which had no unusual repeat annotations. We set *k* = 20 for this sequence, which is similar to what was used in the Mash paper [20]. Less than 1% of the *k*-mers are non-singletons.
2. *D-med*: This is the sequence of RBMY1A1, a chrY gene that is composed of ALUs, SINEs, LINEs, simple repeats, and other repeat elements [Fig 2C in [21]]. We also use *k* = 20 for this sequence. Approximately 3% of *k*-mers are non-singletons.
3. *D-hard*: This is a subsequence of RBMY1A that is annotated as a simple repeat by Tandem Repeats Finder [4], containing 4.2 similar copies of a repeat unit of length 545nt. We use *k* = 10, which is large enough to avoid spurious repeats in this short sequence but small enough to capture its repetitive structure. More than 40% of the *k*-mers are non-singletons.
4. *D-hardest*: This is a subsequence (100k long) of a region that is annotated as ‘Active *α*Sat HOR’ in the chr21 centromere. The location of the subsequence within the region is arbitrary. Alpha satellite (*α*Sat) DNA consists of 171-bp monomers arranged into higher-order repeats, and is notoriously difficult to assemble or map to [17]. We use *k* = 30 for this sequence, as a user dealing with such a sequence is likely to choose a higher *k* value. Over 70% of the *k*-mers are non-singletons.

Before proceeding with experiments, we assess the validity of the two approximations made in the derivation of our estimator. The first approximation is ignoring the dependency between overlapping occurrences of a *k*-mer. The *k*-mers where this happens, i.e. *k*-mers *τ* where *sep*(*τ*) *< occ*(*τ*), contribute to inaccuracy. As shown in Table 1, this is exceedingly rare. The second approximation is ignoring the possibility that a *k*-mer mutates to another *k*-mer in the spectrum. *K*-mer pairs in *s* that have a low Hamming distance will contribute to the bias. Figure 2 shows the distribution of all-vs-all pairwise *k*-mer Hamming distances. The *D-hard* and *D-hardest* datasets indeed have a large amount of “near-repeat” *k*-mers, which should make these datasets challenging for our estimator.

**Figure 2.**
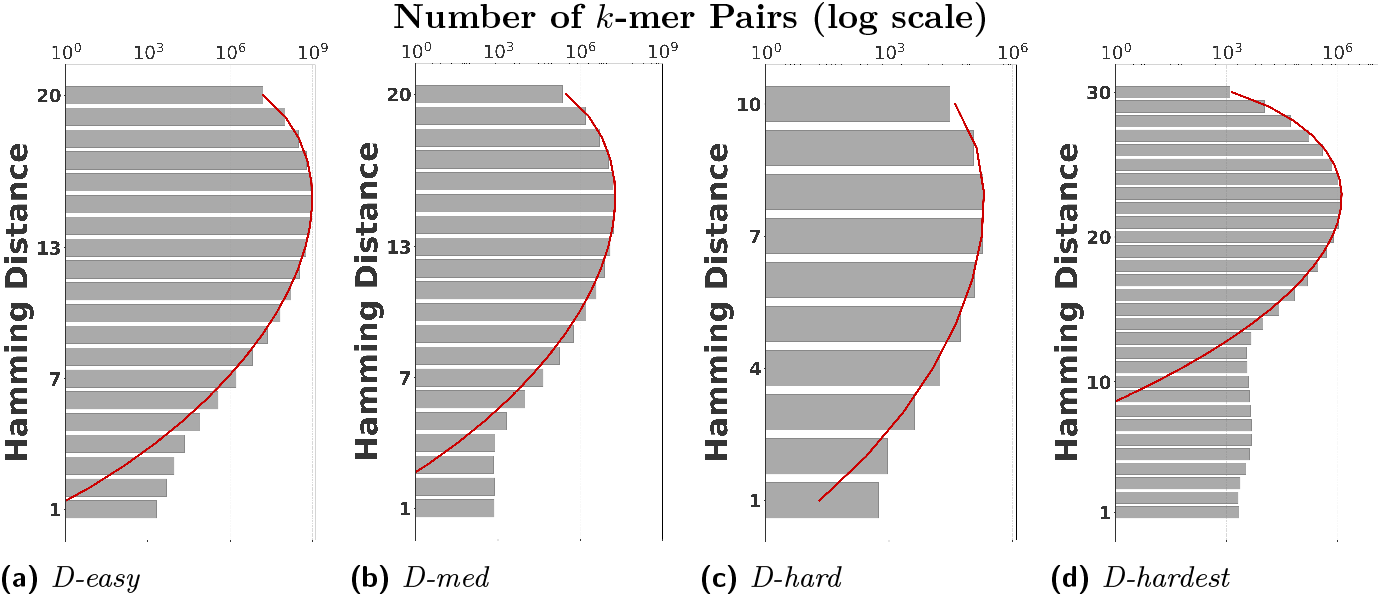
The distribution of all-vs-all *k*-mer Hamming distances. The theoretical Hamming distance distributions between random *k*-mers are shown in the red curves.

### 6.2 Comparison of our estimator to the repeat-oblivious estimator

Figure 3 shows the performance on a range of substitution rates, *r* ∈ (0.1%, 33%). For *D-hard* and *D-hardest*, our estimator has a high accuracy (within a few percent of the true value), in the range of around *r* ∈ (0.1%, 24%). The *r*_obl_ estimator, on the other hand, has a much smaller reliability range, e.g. *r* ∈ ∈ (10%, 24%) in *D-hardest*. For example, when the substitution rate is *r* = 1.1%, the average of *r*_obl_ is 10.4%, while the average of 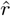 is 1.1%. For *r >* 24%, the observed intersection size was frequently 0; both estimators estimate *r* = 100% at this point, making them unstable. For *D-easy* and *D-med*, the performance of *r*_obl_ is nearly as good as our estimator, except at very low values of *r* (e.g. *r*_obl_ has a 230% relative error at *r* = 0.1% on *D-med*).

**Figure 3.**
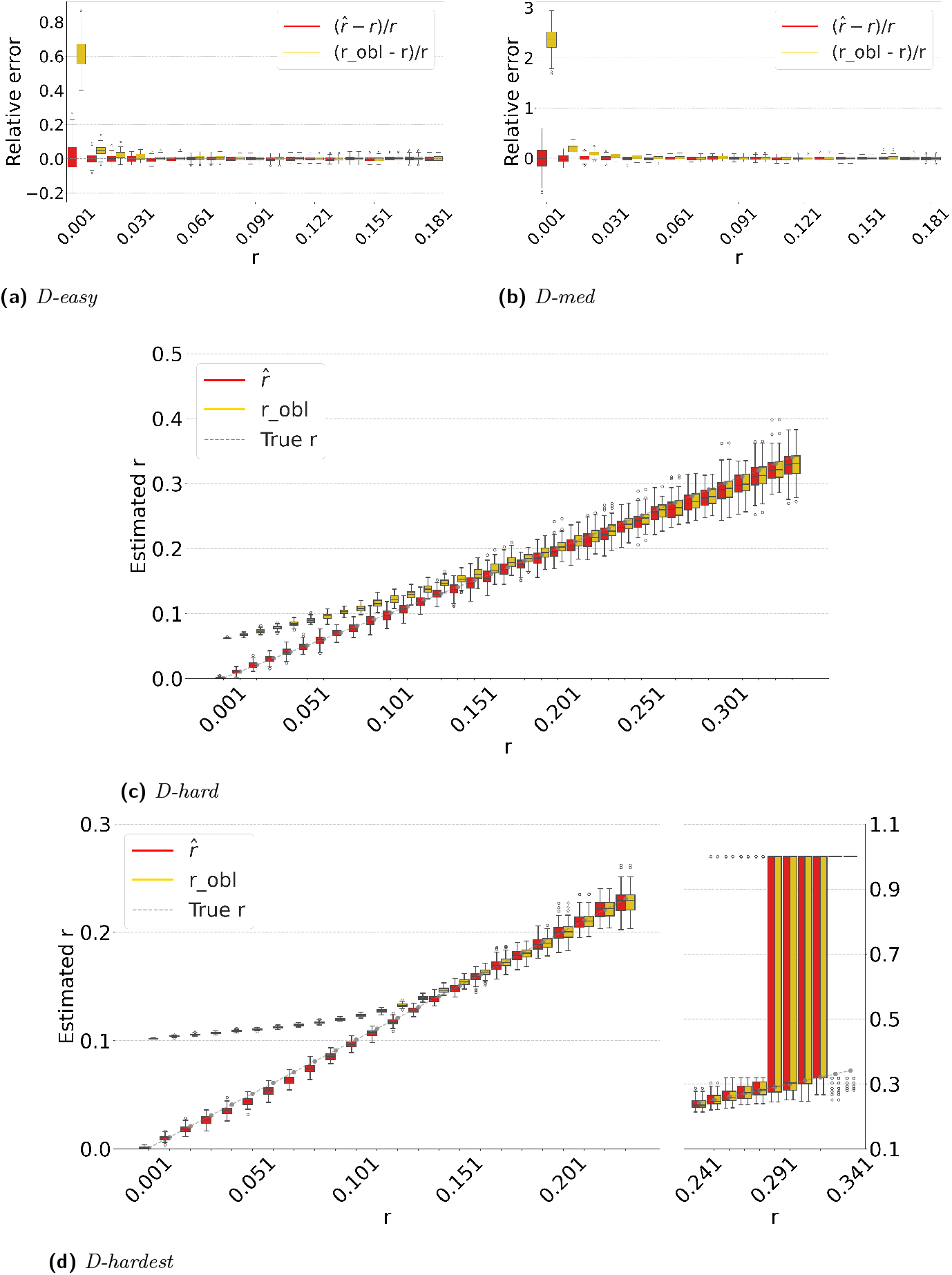
Comparison of our estimator 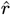 with *r*_obl_. For each *r* value, we simulate the random substitution process 100 times and show the box plot of the resulting estimates. For *D-easy* and *D-med*, the y-axis shows the relative error. For *D-hard* and *D-hardest*, the y-axis shows the actual estimator value instead, in order to reflect the bigger scale of the differences. For *D-easy* and *D-med*, the plots follow the same pattern if they were to be extended rightwards up to *r* = 33%.

Figure 4 evaluates the estimators on *D-hardest* while fixing *r* = 1% and varying *k*. For *k ≥* 690, both estimators become unstable (not shown in figure); similar to the case of high substitution rates, the observed intersection size was frequently 0. For smaller *k*, our estimator performed much better than *r*_obl_, e.g. for *k* = 32, the average *r*_obl_ was 10%.

**Figure 4.**
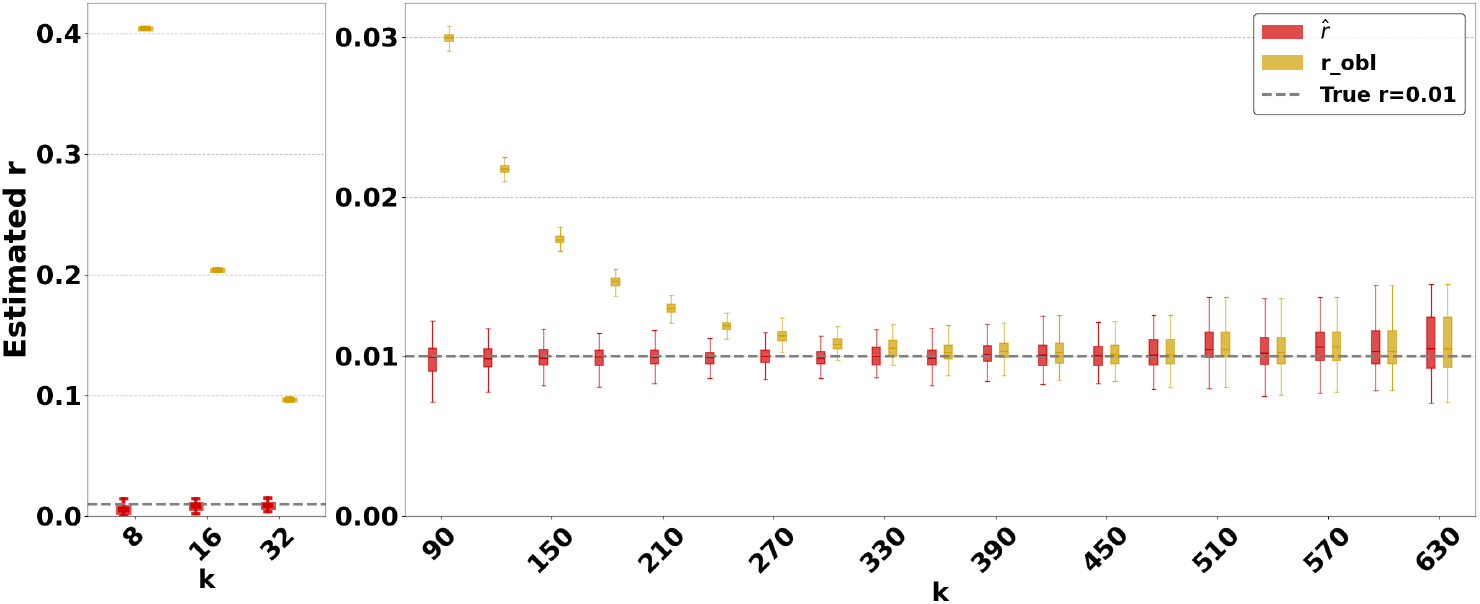
Comparison of our estimator with *r*_obl_ on *D-hardest*. We fixed *r* = 1% and varied *k*. For each *k* value, we simulate the random substitution proess 100 times and show the box plot of the resulting estimators.

The relative performance of the two estimators can be explained algebraically. The *r*_obl_ estimator is derived using the approximation that the probability that a *k*-mer *τ* from *s* remains after substitutions as *occ*(*τ*)(1 − *q*). Our estimator uses the approximation that *τ* remains as 1 − *q*^*occ*(*τ*)^. For singleton *k*-mers, these probabilities are equal, but for repetitive sequences, the effect of *occ*(*τ*) *>* 1 cannot be neglected; therefore, *r*_obl_ gets progressively worse as the datasets become more repetitive. Furthermore, *occ*(*τ, s*)(1 − *q*) *>* 1 − *q*^*occ*(*τ,s*)^ on *q* ∈ (0, 1). Consequently, *r*_obl_ tends to be higher than 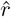. The difference between 1 − *q*^*occ*(*τ,s*)^ and *occ*(*τ, s*)(1 − *q*) increases as *q* decreases. Hence, the gap between *r*_obl_ and 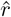 is larger for smaller *r* and smaller *k*, as Figures 3 and 4 show. Finally, as *q* approaches 1, the probability of an empty intersection becomes greater, leading all estimators to output 1. This explains the pattern for large *r* in Figure 3d.

### 6.3 Combination with sketching

Sketching is a powerful technique that can make it possible to quickly compute all-pairs estimates on large datasets [20]. Our estimator lends itself to being applied on the sketched (rather than full) intersection, as follows. Given a threshold 0 *< θ <* 1, one can use a hash function to uniformly map each *k*-mer to a real number in (0, 1). A *FracMinHash sketch* of a sequence *s* is defined as the subset of the *k*-spectrum of *s* that hashes below *θ* [11]. In this way, one can compute *I*^*θ*^, the size of the intersection between the sketches of *s* and *t*.

Recall that our estimator is defined by finding the unique value *r* to solve 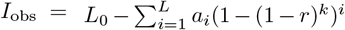, where *I*_obs_ is the size of the observed (non-sketched) intersection. It is easy to show that the expected value of *I*^*θ*^ over the sketching process is *θI*. A natural extension is then to find the unique value of *r* to solve 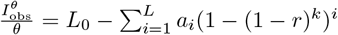. The only caveat is that in some rare cases for very low mutation rates, 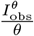 may exceed *L*_0_ and result in a lack of unique solution; in such cases, we hard code the estimator to return 0.

Figure 5a shows the accuracy of the resulting estimator on *D-hardest*, averaged over the combined replicates of the substitution and sketching process. The sketching does not introduce any systematic bias, but, as expected, increases the variance of our estimator. The variance is larger for smaller *θ* values. These results indicate that our estimator can indeed be applied to FracMinHashed sequences, with the threshold parameter *θ* controlling the trade-off between sketch size and the estimator’s variance.

**Figure 5.**
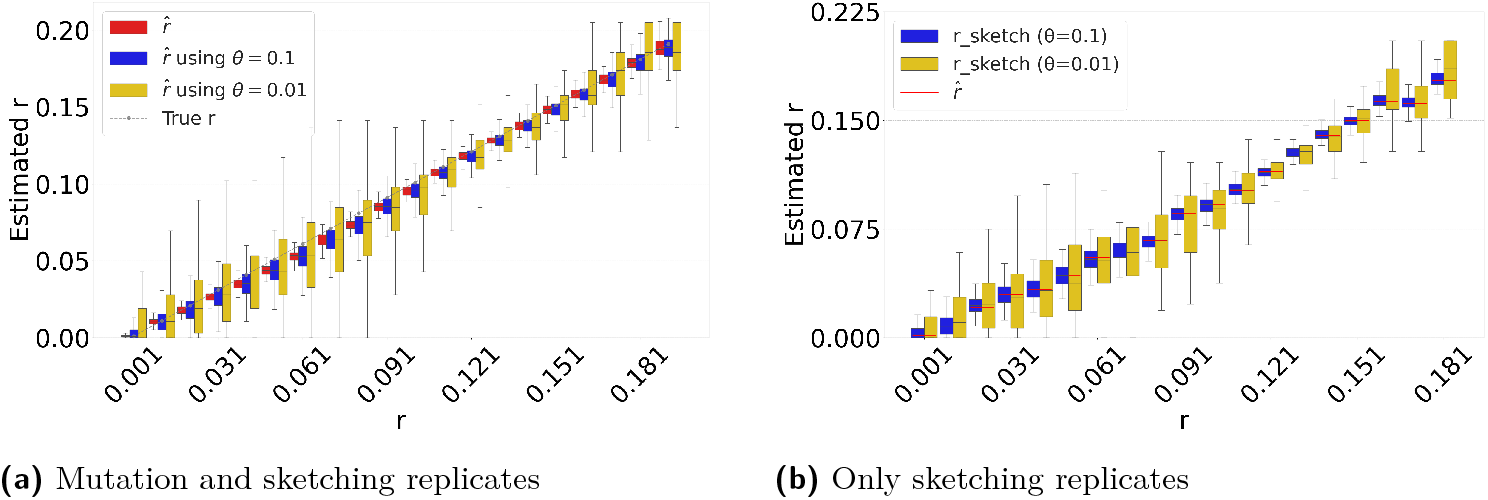
Sketching-based estimation results on *D-hardest*. In panel (a), for each *r*, we replicate the substitution process 100 times and, for each replicate, we replicate the sketching process 100 times. In panel (b), for each *r*, we generate one mutated string and replicate the sketching process 100 times.

Figure 5b evaluates the isolated impact of the sketching process for a fixed string *t*, which better reflects the typical user scenario. For each substitution rate *r*, we generate a single mutated string *t* and compute the 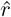 estimate based on the non-sketched intersection. We then replicate the sketching procedure for *s* and *t* and compare the distribution of the sketched estimator to the value of the non-sketched estimate (shown as red bar). The results demonstrate that sketching can accelerate the estimation process, at the cost of introducing controlled variance in the estimates.

### 6.4 Accuracy as a combined function of *k* and *r*

The accuracy of our estimator 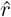 ultimately depends on an intricate interplay between *k* and *r*. A smaller *k* increases the number of repeats, making estimation more challenging. On the other hand, as *r* or *k* increases, the probability *q* = 1 − (1 − *r*)^*k*^ increases, leading to a higher chance of an empty intersection size and an unreliable estimator. To more thoroughly explore the space of all values, Figure 6 evaluates the average relative absolute error, defined as 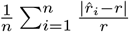, over a wide range of *r* and *k*. This combines our estimator’s empirical bias and variance, indicating the parameter ranges at which our estimator is reliable.

**Figure 6.**
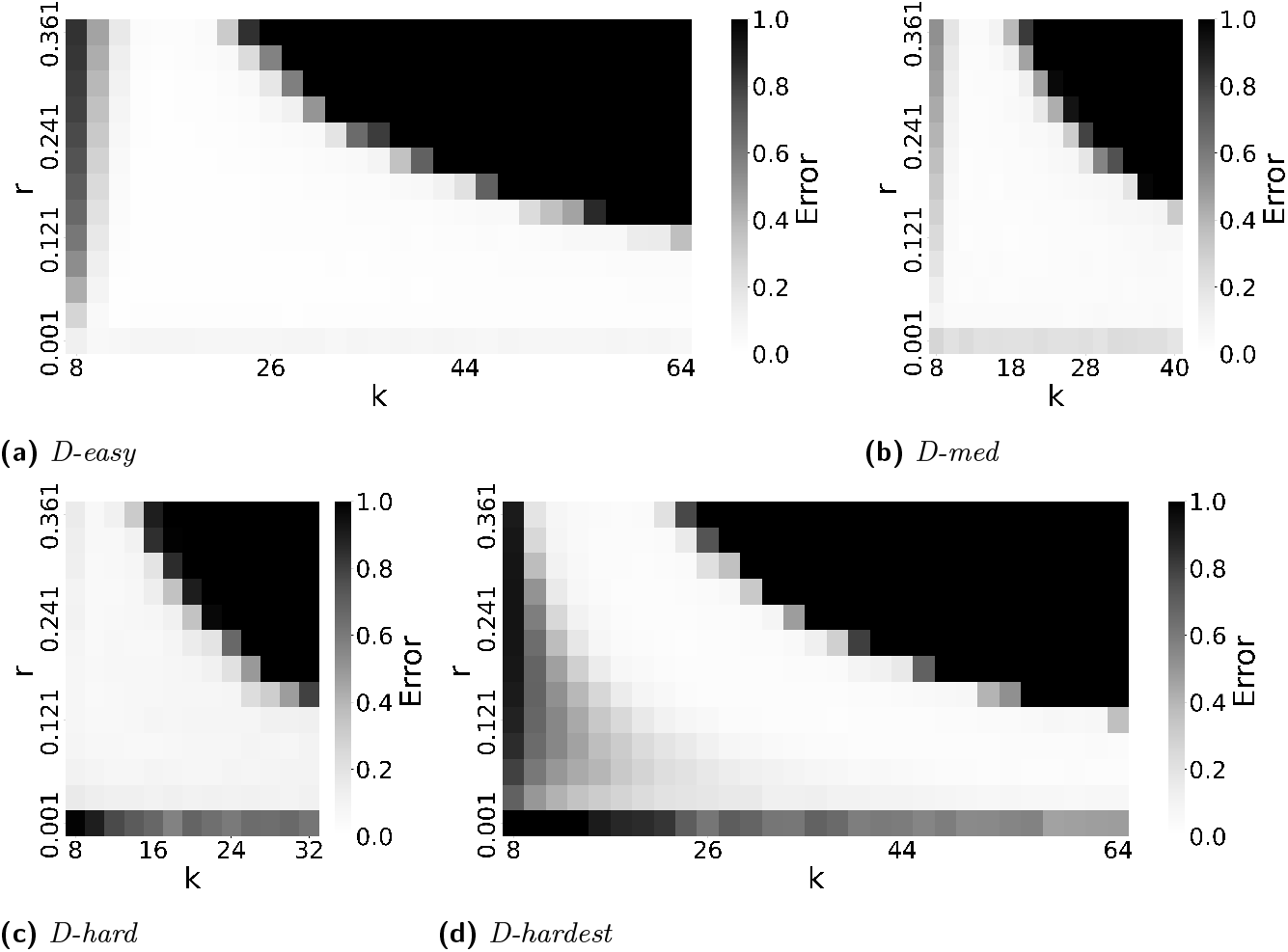
The accuracy of our estimator 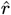 as a function of both *k* and *r*. Each cell shows the average relative absolute error of 100 replicates, e.g. an error of 0.5 means that the estimate is off by 50%. The errors are capped at 1.0, i.e. all errors greater than 1.0 are shown as 1.0.

We note that a user is usually able to choose *k* but not *r*. For *r*, they typically have only a rough range on what it might be. For instance, substitution rates of more than 25% are unlikely for biologically functional sequences. Therefore, choosing a *k* boils down to choosing a column from the heatmap that is good for the desired *r* range. Figure 6 shows that choosing a *k* in the range of 10 to 20 would work well for all of our datasets.

### 6.5 Theoretical bounds on the bias

Theorem 3 gives theoretical bounds on the bias of 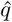. To validate these bounds empirically, we run simulations, using the same setup as in Figure 3. Figure 7 shows that the empirical mean usually lies within the bias bounds, as the theory predicts. In cases where it does not, the empirical variance is high, indicating that the empirical mean has not yet converged to within the bounds. Furthermore, we see that the upper bound is nearly tight. This is consistent with the fact that overlapping *k*-mers are rare (Table 1), implying that that ℱ (*q*) is approximately equal to our lower bound on the expected intersection size (i.e. *L*_E_).

**Figure 7.**
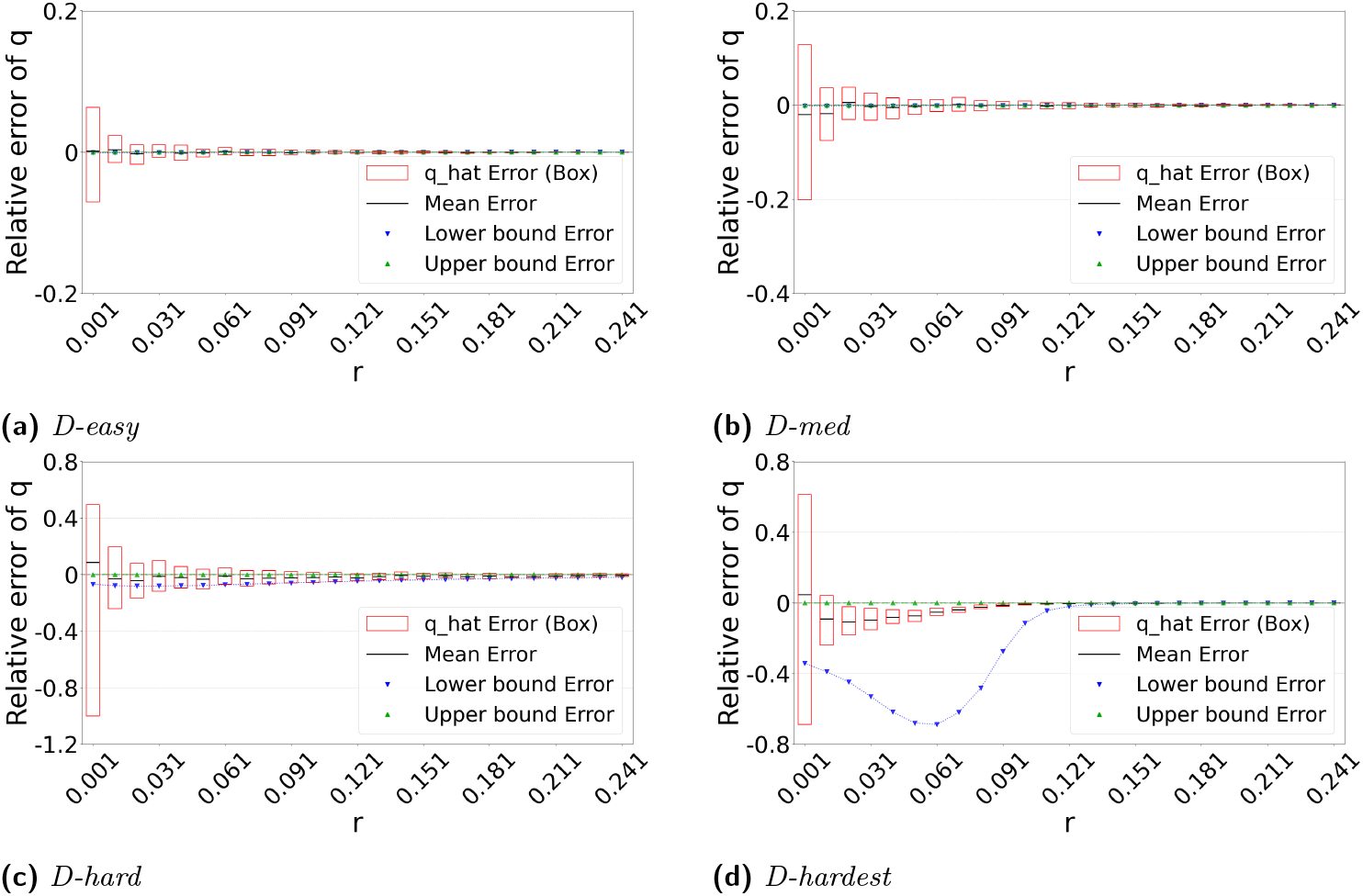
Theoretical bounds on the bias of 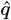. For each *r*, the box plot shows 100 replicates of the substitution process. For the box plots, the y-axis shows the distribution of 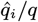. For the lower and upper bound curves, the y-axis corresponds to the ratio of the bound to the true *q*. The black bars in the center of each box represent the mean, rather than the median.

The lower bound tracks the true value closely, except in the range of *r* ∈ (0, 10%) of *D-hardest*. We believe this is primarily due to the looseness of the variance upper bound *U*_*V ar*_ in Theorem 2. When we plugged the observed empirical variance of *I* in place of *U*_*V ar*_ in Theorem 3, the lower bound curve no longer behaved abnormally in *D-hardest* (plot not shown). Furthermore, when we additionally replaced both *U*_E_ and *L*_E_ with the observed empirical mean of *I*, the bounds closely captured the empirical mean of 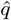. These empirical results suggest that the estimator satisfies the approximation 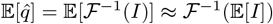. In other words, when we have looseness in the bias bounds, it is due to the looseness of Theorems 1 and 2 rather than Theorem 3.

### 6.6 Identifying unstable parameters using Theorem 4

Figure 6 indicates that when *k* and *r* are large enough to lead to a high *q*, our estimator becomes unstable. Our observations indicate that this happens because the intersection becomes empty, resulting in 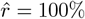 regardless of the true mutation rate. This limitation is anticipated and reflects a fundamental constraint shared by any intersection-based estimator. Figure 7 does not reflect this limitation, because in such cases, the relative error is small simply by virtue of *q* being close to 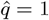 (even though the estimate of *r* is not accurate). We therefore looked for an alternative method to *a priori* determine, given a high value of *k*, which values of high *r* make our estimator unstable.

We hypothesized that computing the probability of an empty intersection size *a priori* can identify such unstable regions of the parameter space, without needing to do simulations as for Figure 6. Though computing this probability is challenging in the general case, Theorem 4 gives an upper bound *P*_empty_ based on only *L, k*, and *r*. The upper bound is approximately tight when not considering the effect of repeats. We therefore hypothesized that when *P*_empty_ is high, our estimator becomes unstable.

Figure 8a plots *P*_empty_ against the accuracy of our estimator. As hypothesized, the substitution rate at which our estimator starts to becomes unstable (around 24 − 28%) coincides with a sharp increase in *P*_empty_. To test this more thoroughly, we computed *P*_empty_ for all values of *k* and *r* for which we evaluated *D-hard* in Figure 6c. Figure 8c shows that there is a close correspondence between *k* and *r* values where our estimator’s relative error is high and *P*_empty_ is high. These observations suggest that *P*_empty_ is a useful diagnostic criterion for determining values of *k*, and *r* when 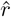 may fail.

**Figure 8.**
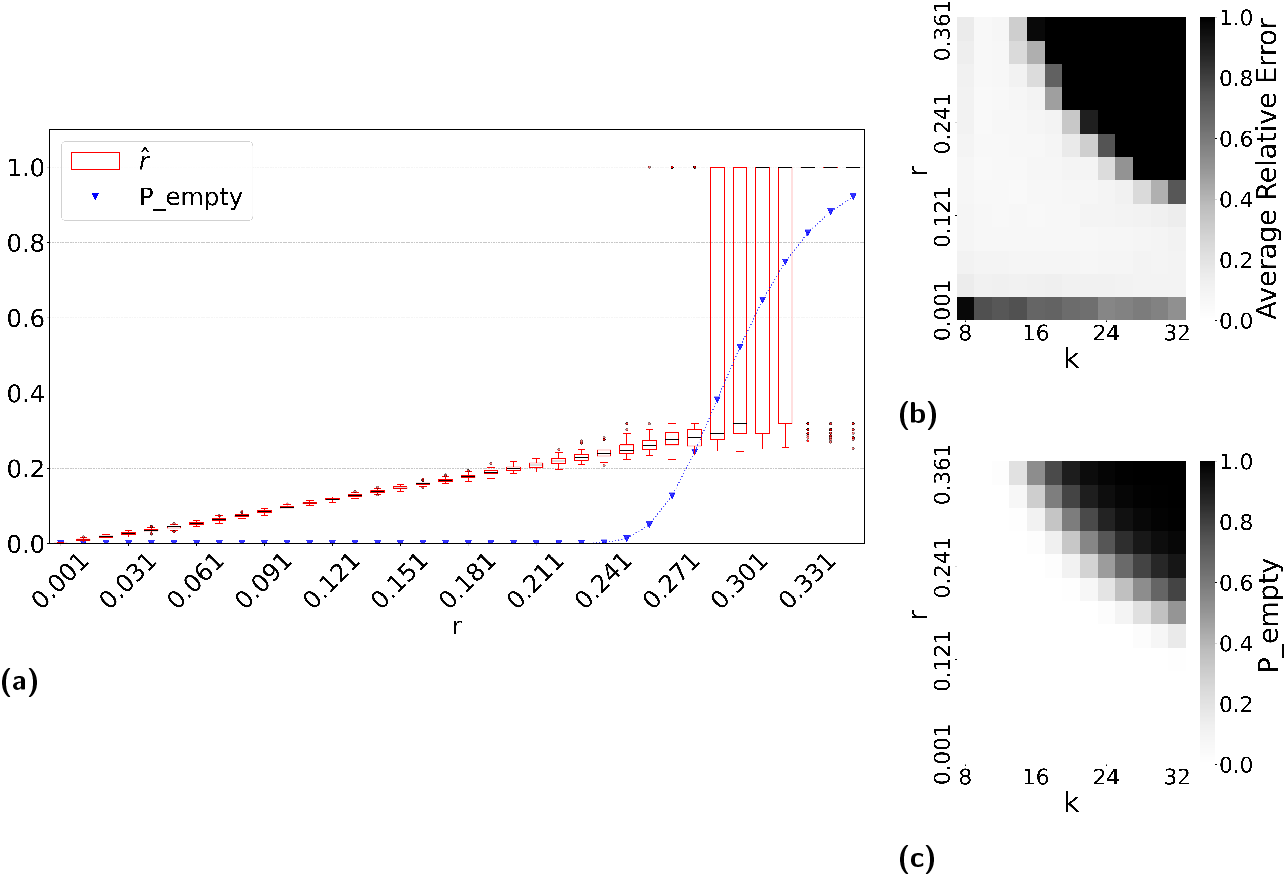
The usefulness of *P*_empty_ as a diagnostic criterion for when our estimator becomes unstable. Panel (a) overlays the estimator values on *D-hardest* with *P*_empty_ values. Panel (b) recapitulates the heatmap of Figure 6c, i.e. the estimator error on *D-hard*. Panel(c) shows the value of *P*_empty_ for the length of *D-hard* and the same parameter values in (*b*).

## 7 Conclusion

In this paper, we propose an estimator for the substitution rate between two sequences that is robust in highly repetitive regions such as centromeres. Our experiments validated its performance across a broad range of *k* and *r* values. We provide theoretical bounds on our estimator’s bias (specifically on the bias of 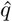), and show that it accurately captures the estimator’s empirical mean in most scenarios.

For large values of *k* and *r*, i.e., when *q* is large, the intersection of the *k*-spectra tends to be empty with high probability, which is a foreseeable limitation for all intersection-based estimators. To address this, we introduce a heuristic criterion, *P*_*empty*_, which depends only on the number of *k*-mers *L*, the *k*-mer size *k*, and the substitution rate *r*. This criterion allows us to heuristically identify parameter settings under which the estimator becomes unstable.

We also showed how our estimator can be easily combined with FracMinHashing. Empirical results show that sketching does not introduce systematic bias, albeit at the cost of increased variance.

We do not perform a runtime analysis of our estimator because it completes in less than a second on our data. The runtime of our estimator is the time it takes to solve an equation numerically using Newton’s method. Since ℱ (*q*) is a polynomial and the solution is constrained to the interval [0, 1], Newton’s method converges in 𝒪 (log log(1*/ϵ*)) iterations, where *ϵ* is the target precision. Each iteration involves evaluating ℱ and its derivative, which takes time proportional to the number of non-zero *a*_*i*_ terms. Except for esoteric corner cases, the number of such terms is small in practice.

The immediate open problem is to tighten the theoretical bounds on the bias. Future work could thus focus on deriving a tighter variance bound to strengthen the theoretical characterization of 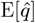. A bigger open question is how to derive confidence intervals. This is a more challenging problem than bounding the bias because it requires a deeper understanding of the estimator’s distribution.

Our estimator could potentially be extended to work on unassembled sequencing reads, as opposed to assembled genomes. Our method does not rely on the *k*-mer multiplicities in the intersection size, making it amenable to such a scenario. Still, one of the limitations of our estimator is the need to know the abundance histogram of the source string. A tool like GenomeScope [31] can estimate the abundance histogram from sequence data *k*-mer counts. Alternatively, the user may choose to use an abundance histogram from a related genome, as related genomes are likely to have similar abundance histograms. Fully adapting this estimator to work with sequencing data remains an important future work.

## Acknowledgements

We thank Amatur Rahman for initial work on the project and Qunhua Li and David Koslicki for helpful discussions. We thank Mahmudur Rahman for the helpful discussion about hash functions. We thank Bob Harris for the idea of using dynamic programming to compute the probability of the destruction of all k-spans.

## A Appendix: Missing Proofs

### Theorem 4.

**Proof**. Let *M*_*i*_ be the event that every interval of length *k* in *s*[1 … *i* + *k* − 1] has at least one substitution. We claim that the following recurrence holds, which automatically leads to the dynamic programming algorithm of the desired time.

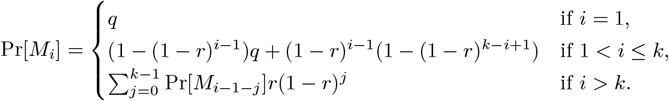

We will use *k-span* to denote an interval of length *k* in *s*. For the case that *i* = 1, *M*_*i*_ is the probability that the first *k*-mer mutates, which is *q*. For 1 *< i ≤ k*, we do the following. We denote *A* as the event that there is at least one substitution in *s*[1, *i* − 1], *B* as the event that there is at least one substitution in *s*[*i, k*], and *C* as the event that there is at least one substitution in *s*[*k* + 1, *i* + *k* − 1]. Then we have

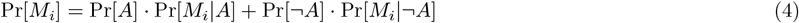

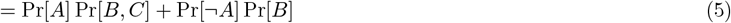

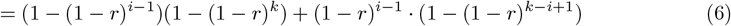

For the last case (*i > k*), let *L*_*j*_ be the event that *j* is the position of the rightmost substitution in *s*[1, *i* + *k* − 1]. Observe that for *j* ≠ *ℓ, L*_*j*_ and *L*_*ℓ*_ are mutually exclusive. Furthermore, observe that the rightmost mutation position must be at least *i*, otherwise the *k*-span starting at *i* is not mutated. Therefore, by the law of total probability, we have

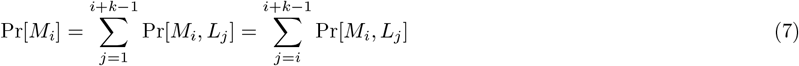

Observe that if there is a substitution at position *j*, then all the *k*-spans beginning at positions *j* − *k* + 1, …, *i* are mutated. Therefore, Pr[*M*_*i*_, *L*_*j*_] = Pr[*M*_*j* − *k*_, *L*_*j*_] = Pr[*M*_*j* − *k*_] *·* Pr[*L*_*j*_]. Hence,

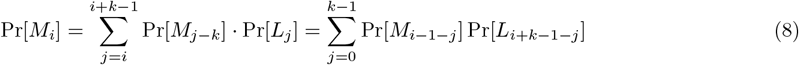

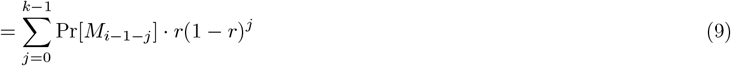

□

### Lemma A.1.

*Let X be a sum of random variables X*_1_, …, *X*_*n*_, *and let µ* = E[*X*]. *Then*

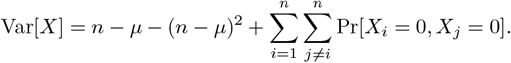

**Proof**.

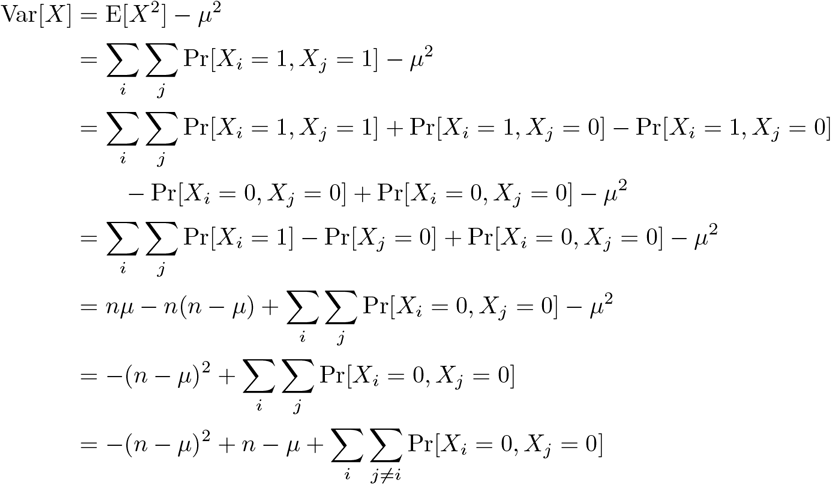

▪

### Lemma A.2.

ℱ (*q*) *is invertible on* [0, 1]. *Moreover, if there exists at least one k-mer τ with occ*(*τ*) = 1, *then, on the intervals q* ∈ [0, 1] *and y* ∈ [ℱ (1), ℱ (0)], *and letting f denote the inverse of* ℱ, *we have that*

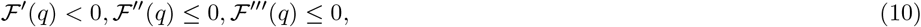

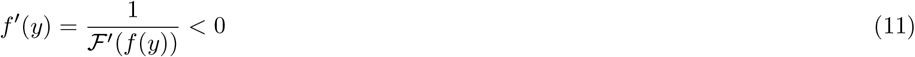

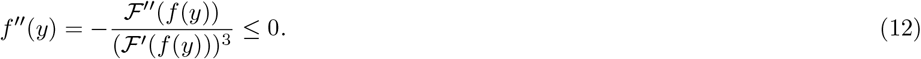

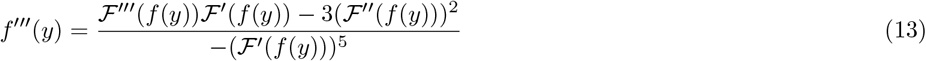

**Proof**. Recall that *a*_*i*_ is the number of *k*-mers that have *i* copies, and that 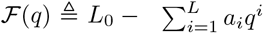. Since all *a*_*i*_ values are non-negative, all derivatives of ℱ are non-positive on *q* ∈ [0, 1]. Moreover, since we assume that *a*_1_ is strictly positive, the first derivative of ℱ is strictly negative on *q* ∈ [0, 1]. The derivatives of *f* can be expressed in terms of the derivatives of ℱ and *f* by applying the inverse function rule from basic calculus.▪

## Footnotes

1 What we describe is based on the estimators used in [8, 20, 12], but with two important differences. The first is that we use the modification adopted in the follow up work of [24] and described in Appendix A.6 of [3]. The second is that the original estimator was calculated from the Jaccard similarity between two sequences; however, under our substitution process model, we can state more simply in terms of the spectrum intersection size.

2 We note that this assumption is not theoretically precise, because forbidding a *k*-mer at position *i* from mutating to *τ* ∈ *s* usually implies that there is at least one *ν* ∉ *s* that the *k*-mer at position *i* + 1 can no longer mutate to. Because of these dependencies, there are downstream effects on the probability space that are complex to track. A theoretically robust alternative was given in [5] via the *k*-span formulation of the problem. It could be used to formalize the assumption here, however, in this paper, we only use the assumption to give intuition for the estimator and do not use it in any formal theorem statements or proofs.

